# MHC-II constrains the natural neutralizing antibody response to the SARS-CoV-2 spike RBM in humans

**DOI:** 10.1101/2020.12.26.424449

**Authors:** Andrea Castro, Kivilcim Ozturk, Maurizio Zanetti, Hannah Carter

## Abstract

SARS-CoV-2 antibodies develop within two weeks of infection, but wane relatively rapidly post-infection, raising concerns about whether antibody responses will provide protection upon re-exposure. Here we revisit T-B cooperation as a prerequisite for effective and durable neutralizing antibody responses centered on a mutationally constrained RBM B cell epitope. T-B cooperation requires co-processing of B and T cell epitopes by the same B cell and is subject to MHC-II restriction. We evaluated MHC-II constraints relevant to the neutralizing antibody response to a mutationally-constrained B cell epitope in the receptor binding motif (RBM) of the spike protein. Examining common MHC-II alleles, we found that peptides surrounding this key B cell epitope are predicted to bind poorly, suggesting a lack MHC-II support in T-B cooperation, impacting generation of high-potency neutralizing antibodies in the general population. Additionally, we found that multiple microbial peptides had potential for RBM cross-reactivity, supporting previous exposures as a possible source of T cell memory.

**Graphical abstract:** 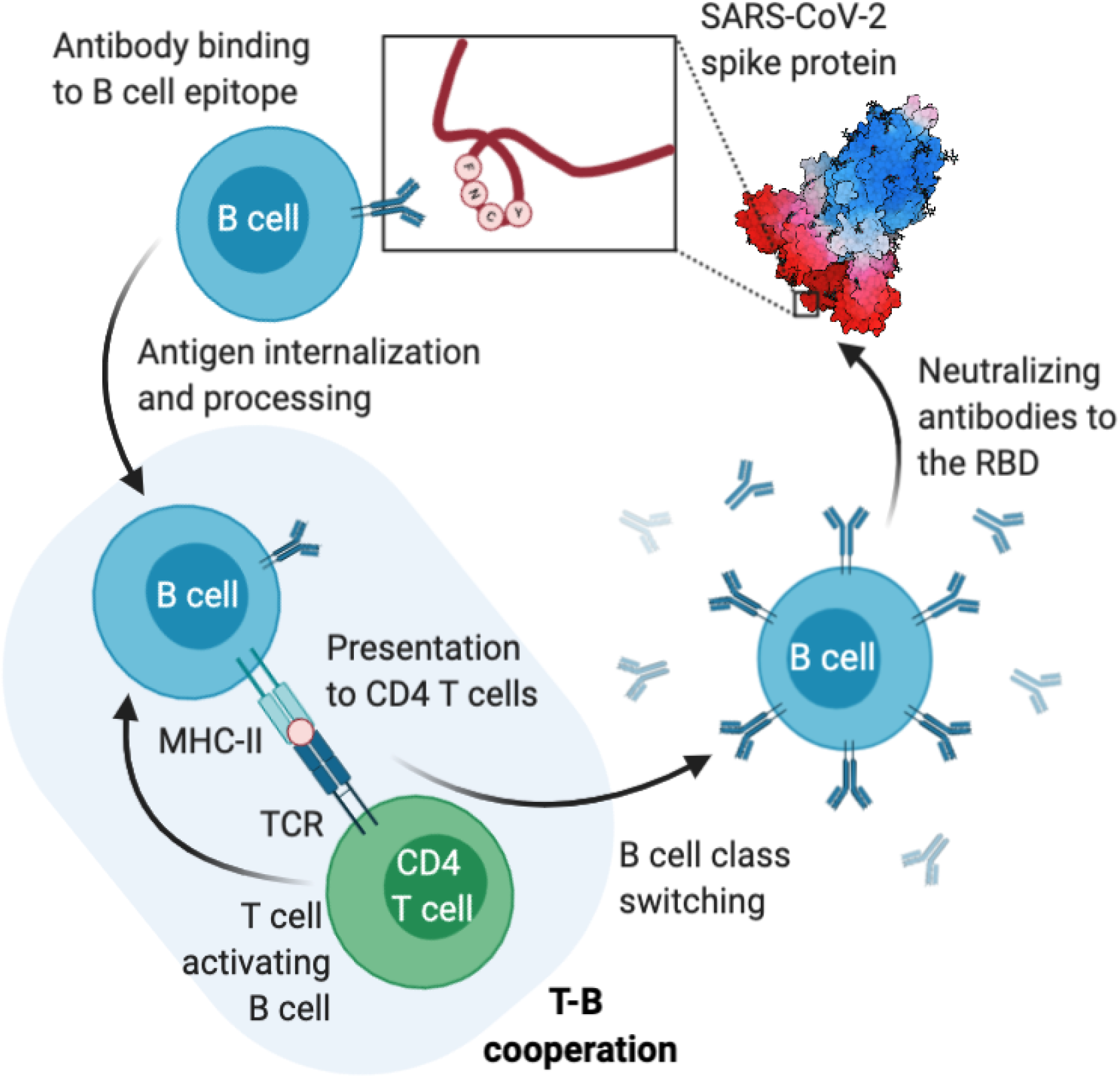

## Introduction

Upon infection with SARS-CoV-2 the individual undergoes seroconversion. In mildly symptomatic patients, seroconversion occurs between day 7 and 14, includes IgM and IgG, and outlasts virus detection with generally higher IgG levels in symptomatic than asymptomatic groups in the early convalescent phase (1). Alarmingly, the IgG levels in both asymptomatic and symptomatic patients decline during the early convalescent phase, with a median decrease of ~75% within 2–3 months after infection (2). This suggests that the systemic antibody response which follows natural infection with SARS-CoV-2 is rapid but short-lived, with the possibility of no residual immunity after 6-12 months (3) affecting primarily neutralizing antibodies in plasma (4).

The generation of an antibody response requires cooperation between a B cell producing specific antibody molecules and a CD4 T cell (helper cell) activated by an epitope on the same antigen as that recognized by the B cell (T-B cooperation) (5). This reaction occurs in the germinal center (6,7). Excluded from this rule are responses against carbohydrates and antigens with repeating motifs that alone cross-link the B cell antigen receptor leading to B cell activation (8). Discovered over 50 years ago (9–11), it also became apparent that T-B cooperation is restricted by Major Histocompatibility Complex class II (MHC-II) molecules (12–14). T-B cooperation plays a key role in the facilitation and strength of the antibody response (10,15) and the size of the antibody response is proportional to the number of Th cells activated by the B cell during T-B cooperation (13,14,16). The importance of T cell help during the activation of antigen specific B cells to protein antigens driving B cell selection is emphasized by recent experiments where the injection of a conjugate of antigen (OVA) linked with an anti-DEC205 antibody induced a greater proliferation of DEC205+ relative to DEC205-B cells consistent with a T helper effect on B cell activation (17).

T-B cooperation requires that the epitopes recognized by the B and T cell be on the same portion of the antigen (11,18,19) leading to a model requiring the contextual internalization and co-processing of T and B cell epitopes (5) which is consistent with the principle of linked (aka associative) recognition of antigen (20). Studies *in vitro* using human T and B lymphocytes showed that an antigen specific B cell can present antigen to CD4 T cells even if antigen is present at very low concentration (10^−11^ – 10^−12^ M) (21). Presentation of antigen by the B cell also facilitates the cooperation between CD4 T cells of different specificities resulting in enhanced generation of memory CD4 T cells (22). However, T-B cooperation is not the only form of cooperative interaction among lymphocytes as cooperation exists between CD4 T and CD8 T cells (23) and between two CD4 T cells responding to distinct epitopes on the same antigen (24).

A model based on coprocessing of T and B epitopes also led to the suggestion that preferential T-B pairing could be based on topological proximity (25–29) so that during BCR-mediated internalization the T cell epitope is protected by the paratope of the BCR. Indeed, a more recent study showed that not only is CD4 T cell help a limiting factor in the development of antibodies to smallpox (vaccinia virus), but that there also exists a deterministic epitope linkage of specificities in T-B cooperation against this viral pathogen (30). Collectively, it appears that T-B pairing and MHC-II restriction are key events in the selection of the antibody response to pathogens and that operationally T-B cooperation and MHC-II restriction are key events in the generation of an adaptive antibody response, suggesting that lack of or defective T-B preferential pairing could result in an antibody response that is suboptimal, short-lived, or both.

In SARS-CoV-2, neutralizing antibodies (NAbs) are a key defense mechanism against infection and transmission. NAbs generated by single memory B cell VH/VL cloning from convalescent COVID-19 patients have been extremely useful in defining the fine epitope specificity of the antibody response in COVID-19 individuals. At present, SARS-CoV-2 NAbs can be distinguished into three large categories. 1) Repurposed antibodies, that is, NAbs discovered and characterized in the context of SARS-CoV and subsequently found to neutralize SARS-CoV-2 via cross-reactivity. These antibodies map away from the receptor binding domain (RBD) of the spike protein (31–33). 2) Non-RBD neutralizing antibodies discovered in SARS-CoV-2 patients whose paratope is specific for sites outside the RBD (34). 3) RBD antibodies, including NAbs, derived from SARS-CoV-2 patients that map to a restricted site in the RBD (35–41). Cryo-EM of this third antibody category shows that they bind to residues in or around the four amino acids Phe-Asp-Cys-Tyr (FNCY) in the receptor binding motif (RBM) (residues 437-508) which is inside the larger RBD (residues 319-541) at the virus:ACE2 interface (36). Although the RBD has been shown to be an immunodominant target of serum antibodies in COVID-19 patients (42), high potency NAbs are directed against a conserved portion of the RBM on or around the FNCY patch, a sequence only found in the RBD of SARS-CoV-2 and not in other coronaviruses. Indeed while the RBD is mutationally tolerant, the RBM is constrained to the wild-type amino acids (43), implying that the B cell epitope included in this region of the virus:ACE2 interface is resistant to antigenic drift. Thus, we may refer to this site as a key RBM B cell epitope in the generation of potent NAbs.

Antibody responses against SARS-CoV-2 depend on CD4 T cell help. Spike-specific CD4 T cell responses have been found to correlate with the magnitude of the anti-RBD IgG response whereas non-spike CD4 T cell responses do not (44). However, spike-specific CD4 T cells reactive with MHC-II peptides proximal to the central B cell epitope represent a minority (~10%) of the total CD4 T cell responses, which are dominated by responses against either the distal portion of the spike protein or other structural antigens (45). Surprisingly, these CD4 T cell responses are largely cross-reactive and originate from previous coronavirus infections (46).

As mounting evidence suggests that the NAb response in COVID-19 patients is relatively short-lived, we decided to test the hypothesis that associative recognition of the key RBM B cell epitope and proximal MHC-II-restricted epitopes may be defective with detrimental effects on preferential T-B pairing. Therefore, to quantify the potential effects of T-B cooperation *in vivo*, we analyzed all 15mer putative MHC-II epitopes (+/− 50 amino acid residues) relative to the key RBM B cell epitope for coverage by all known 5,620 human MHC-II alleles and predicted binding affinity. The analysis shows that there exists in general less availability of effective T cell epitopes in close proximity to the key RBM B cell epitope in the human population.

## Results

### Topology of a key RBM B cell epitope

Within the 222 amino acid long RBD of the spike protein (residues 319-541), the RBM (residues 437-508) is the portion of the spike protein that establishes contact with the ACE2 receptor (**Fig 1A**). The contact residues span a relatively large surface involving approximately 17 residues (36), among them residues F486, N487, Y489 form a loop, which we term the FNCY patch, which is surface exposed and protrudes up towards the ACE2 receptor from the bulge of the RBD (**Fig 1B-C**). F486 forms hydrophobic interactions with three ACE2 residues (L79, M82, W83). N487 forms hydrogen bonds with Q24 and W83, and Y489 is linked with K31 via a hydrophobic interaction. This makes the amino acid residues in or around the FNCY patch a logical B cell epitope target for antibodies blocking the virus:receptor interaction. In addition, these core residues are mutationally constrained by the ACE2 contact surface (43). Not surprisingly, a set of recently reported potently neutralizing antibodies generated by single B cell VH/VL cloning from convalescent COVID-19 patients all bear paratopes that include the FNCY patch in their recognition site (34,39–41,47) (**Fig 1D**). While other residues (Q493, N501, and Y505) are also shared between ACE2 and the paratope of these antibodies, they are not as protruding and are on a β-sheet unlike the FNCY patch which is organized in a short loop as a result of the C480:C488 disulfide bond. Thus, blockade of the RBM:ACE2 interaction (neutralization) depends at least in part on a B cell epitope in the RBM that is structurally and functionally critical to the interaction, virus internalization, and cell infectivity.

**Figure 1:**
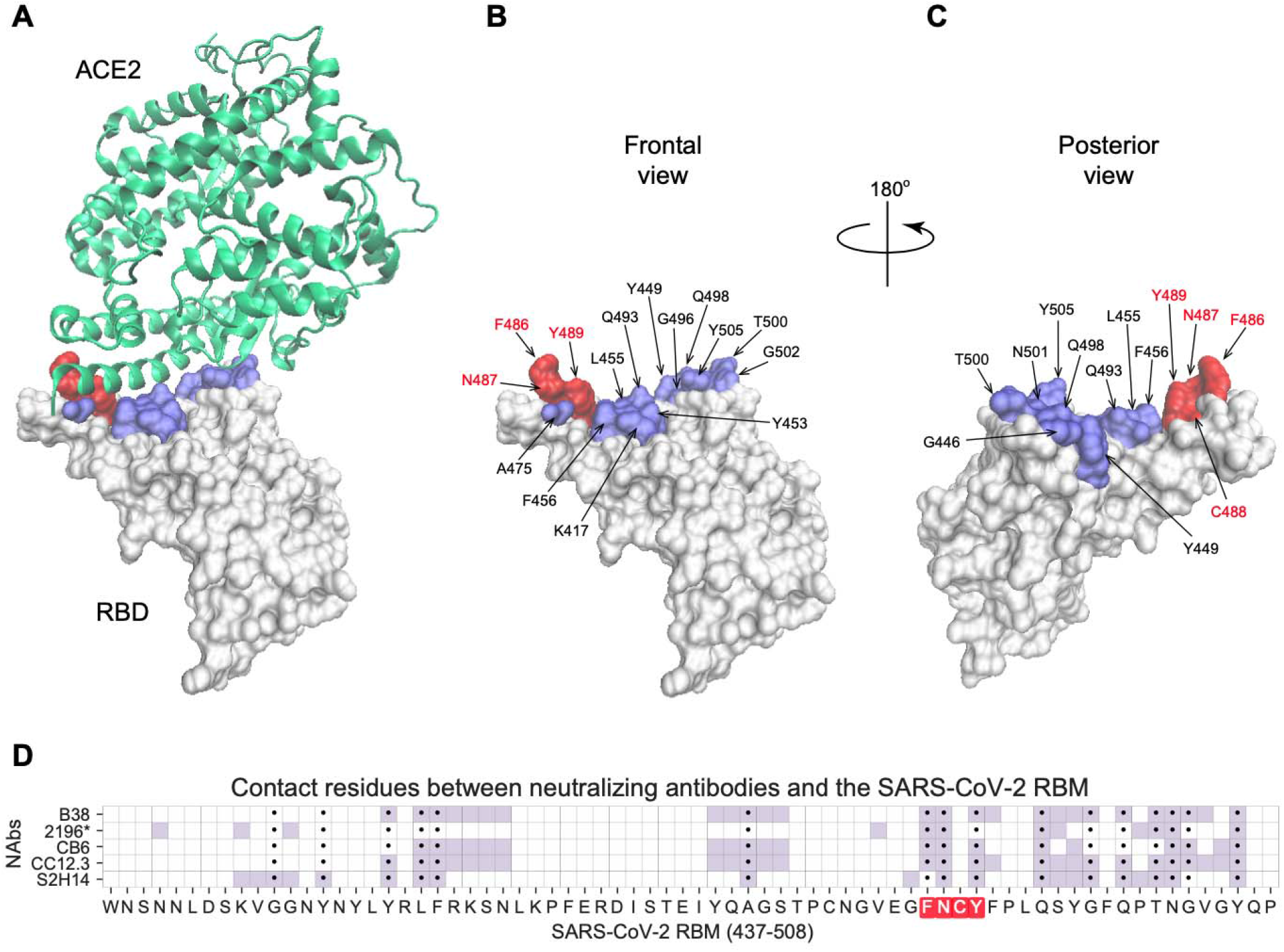
Visualization of the FNCY core of the RBM B cell epitope on the SARS-CoV-2 spike protein RBD. (A) 3D structure of the SARS-CoV-2 spike protein RBD (white) binding the ACE2 receptor (green) (PDB: 6M0J) with contact residues highlighted in blue and the FNCY patch highlighted in red. (B-C) Spike protein RBD with ACE2 contact residues and FNCY patch residues labeled in two orientations (front and back). (D) Heatmap of neutralizing antibody contact residues (purple) on the spik protein RBM region (positions 437-508). Black dots indicate ACE2 contact residues and the FNCY patch is highlighted in red. Source data available in Supplemental Table 1.

### Prediction of MHC-II affinity for 15mer peptides proximal to the RBM B cell epitope

In the T-B cooperation model, B cell activation and production of NAbs is dependent on CD4 T cell responses to MHC-II restricted peptides. To test the hypothesis that the generation of NAbs against a mutationally constrained B cell epitope in the RBM reflects the efficiency of processing and presentation of MHC-II peptides proximal to the FNCY patch, we evaluated the landscape of MHC-II peptide restriction across the entire SARS-CoV-2 spike protein with respect to common MHC-II alleles in the human population. To assess the potential for effective restriction by MHC-II molecules in a reasonable proportion of the population, we devised a position-based score that assigns each amino acid residue the median affinity of the best overlapping peptide, where median affinity is calculated across the 1911 most common MHC-II alleles (**Fig 2A**), which was highly correlated with scores across all 5620 MHC-II alleles (**Fig 2B**; Pearson rho=0.99, p<2.2e-308). While a number of sites along the spike protein are predicted to generate high affinity peptides for most common MHC-II alleles, the region around the FNCY patch was depleted for generally effective binders (**Fig 2C**, Fisher’s exact OR=0.21, p=0.015, Methods, **Supplemental Fig 1**). Interestingly, the RBM region containing the FNCY patch was free of glycans that could potentially mask the epitope (**Fig 2D**). We further evaluated the distributions of binding affinities for the 20 best-ranked peptides across all sites in the spike protein (**Fig 2E**), and in comparison, the distributions for the best 20 peptides overlapping positions within +/− 50 residues of the FNCY patch (**Fig 2F**). In the best case, less than half of the considered MHC-II alleles bound a shared peptide close to the FNCY patch, whereas at other sites there were multiple peptides that could be bound by nearly all of the MHC-II alleles (**Fig 2E**). This suggested overall less availability of effective T cell epitopes in close proximity to the FNCY B cell epitope, which could limit the availability of T cell help during an epitope-specific T-B cooperative interaction in the germinal center.

**Figure 2:**
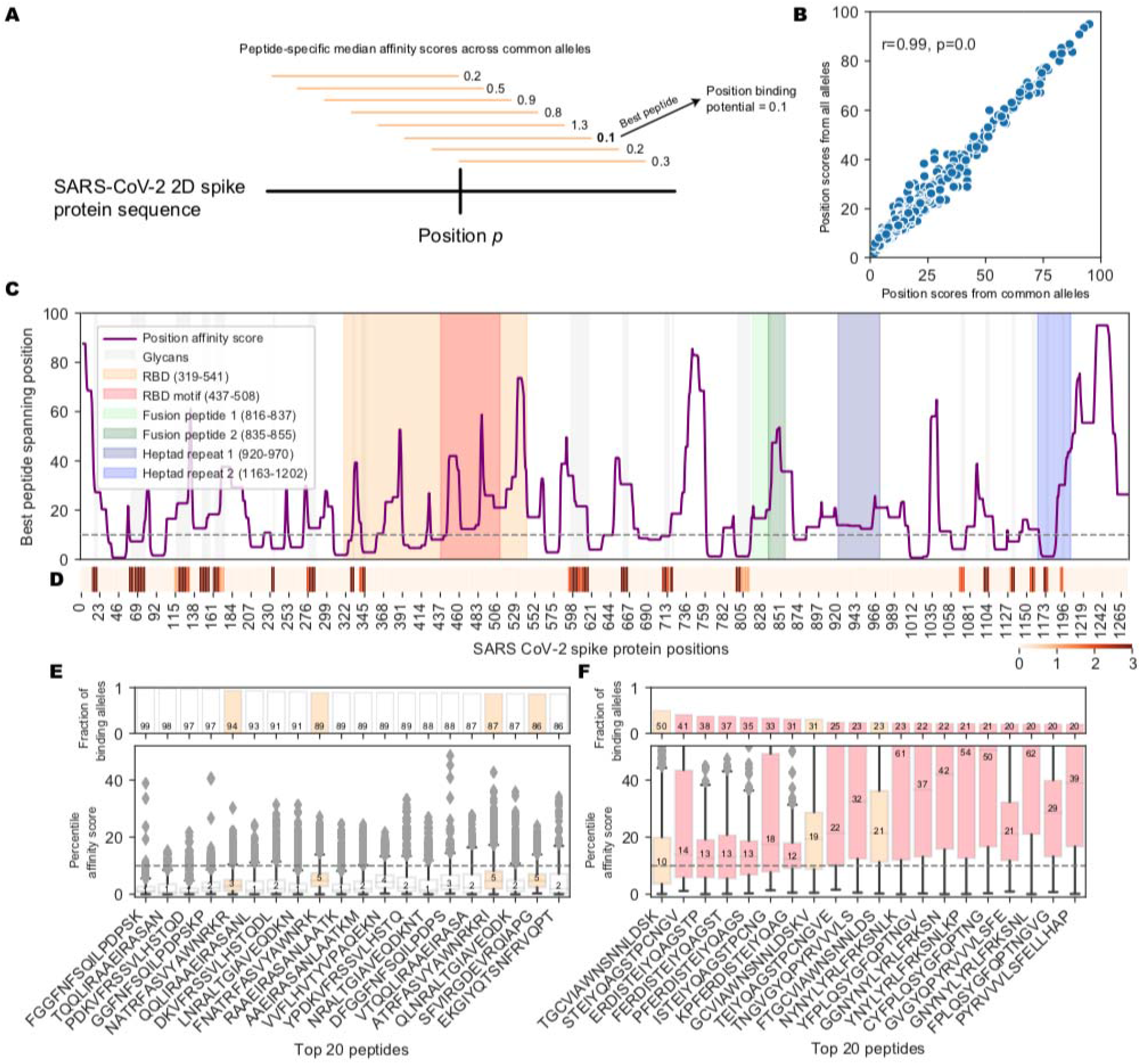
Landscape of MHC-II binding affinity across spike protein 2D sequence. (A) Overview of the position affinity score. (B) Scatterplot showing position affinity scores estimated using only common (>10% frequency, from (48)) MHC-II alleles (x-axis) versus across all MHC-II alleles (y-axis). (C) Lineplot showing the position affinity scores across common MHC-II alleles (Methods). Annotated domains from UniProt are highlighted. (D) Heatmap showing amino acid positions that are glycosylated (49). (E) Barplots (top) and boxplots (bottom) describing the fraction of binding MHC-II alleles and corresponding affinity percentile rank distributions respectively for the top 20 peptides with the highest fraction of common binding alleles. The binding threshold of 10 is shown as a dotted line, with values less than 10 indicating binding. Colors correspond to the regions listed in C. (F) Barplots (top) and boxplots (bottom) describing the fraction of binding MHC-II alleles and corresponding affinity percentile rank distributions respectively for the top 20 peptides within +/− 50 amino acids of the FNCY B cell epitope. Colors correspond to the regions listed in C.

To further assess whether population variation in MHC-II MHC alleles might contribute to heterogeneity in potential to generate neutralizing antibodies, we also evaluated the potential of MHC-II supertypes to restrict peptides from neighboring the FNCY patch. Greenbaum *et al.* previously defined 7 supertypes that group MHC-II alleles based on shared binding repertoire. These 7 supertypes account for between 46%-77% of haplotypes and cover over 98% of individuals when all four loci are considered together (50). We revisited our analysis of peptide restriction proximal to the FNCY patch treating each supertype separately. There was considerable variability in potential to effectively present FNCY patch proximal sequences across supertypes (**Fig 3A-B**, Χ^2^=175, p=3.75e-35, **Supplemental Fig 2**). Only 3 supertypes (DP2, main DP and DR4) commonly presented peptides overlapping the FNCY patch (**Fig 3B**). We were able to obtain population allele frequencies for four populations from the Be The Match registry (51) and Du *et al.* (52). These data show that DR4 is relatively infrequent across the populations evaluated, whereas main DR, main DP, and DP2 are more common (**Fig 3C**), and thus could be more important for MHC-II restriction supportive of neutralizing antibodies. While there were some large population-specific differences in main DP and DP2 supertype frequencies, these frequency estimates are based on a limited population sample and may provide only a rough approximation. In general, DP and DR haplotypes were able to restrict more FNCY patch proximal sequences (**Fig 3D**).

**Figure 3:**
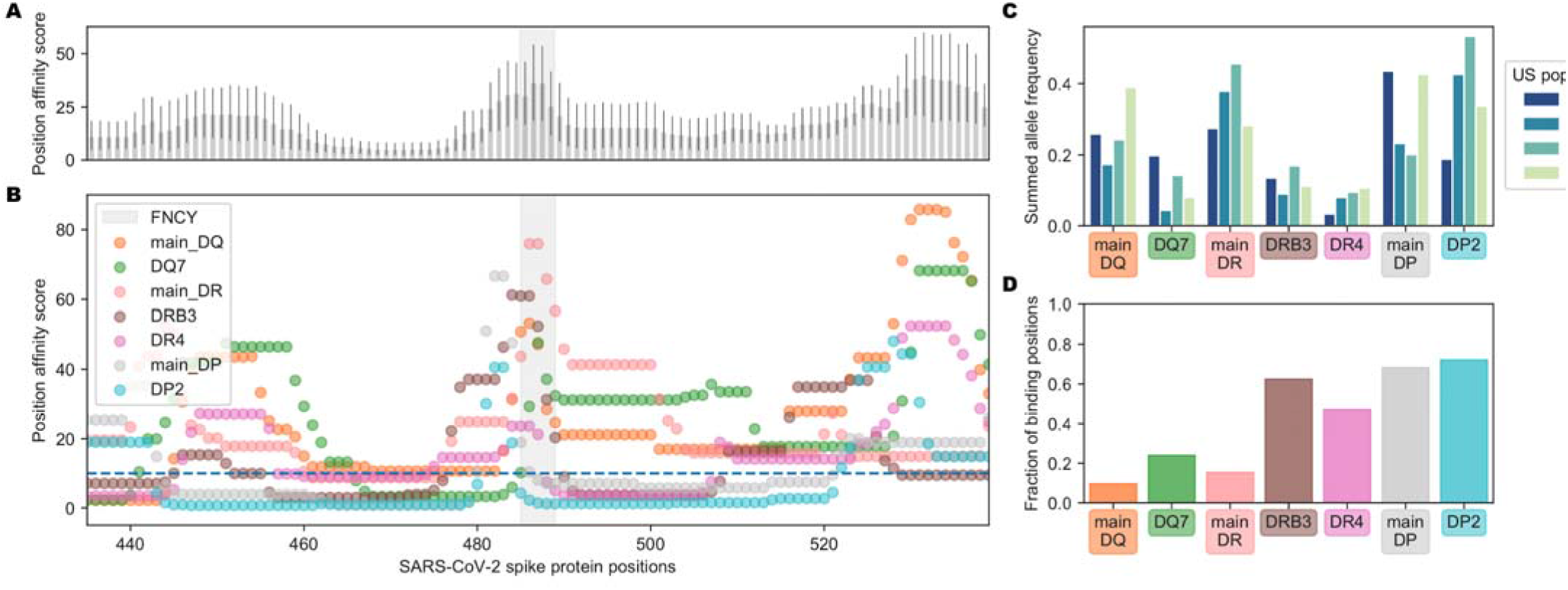
Population variation affecting availability of FNCY proximal T cell epitopes. (A) Barplot showing the aggregated supertype position affinity scores for each position +/− 50 amino acids from the FNCY patch (grey zone). (B) Scatterplot showing the specific supertype position scores for each position +/− 50 amino acids from the FNCY patch (grey zone). The binding threshold of 10 is shown as a dashed blue line, with points below the threshold indicating binding. (C) Barplot showing United States population frequencies, summed across the available alleles in each supertype. (D) Fraction of positions falling below the binding threshold within the region of interest for each supertype.

### Cross-reactivity to a non-coronavirus MHC-II binding peptide as a potential driver of T cell responses helping antibody response to the RBM B cell epitope

Interestingly, Mateus *et al.* reported pre-existing CD4 T cell responses to peptides derived from the spike protein using T cells from unexposed individuals, suggesting previous exposures to other human coronaviruses could potentially generate protective immunity toward SARS-CoV-2. Indeed, regions of higher coronavirus homology were associated with more T cell responses in their data (46). This represents the most comprehensive interrogation of the spike protein with response to CD4 T cell responses to date. They screened all 15mers of the spike protein in pooled format and further evaluated 66 predicted MHC-II peptides that generated CD4 T cell responses. Visualizing the landscape of the CD4 T cell responses described in their work by percent positive response (**Fig 4A**) or spot forming cells (**Fig 4B**), we noted relatively few responses proximal to the FNCY patch in the RBM. Accordingly, few other coronaviruses had limited homology to the FNCY region, and none fully included the FNCY patch (**Fig 5A**).

**Figure 4.**
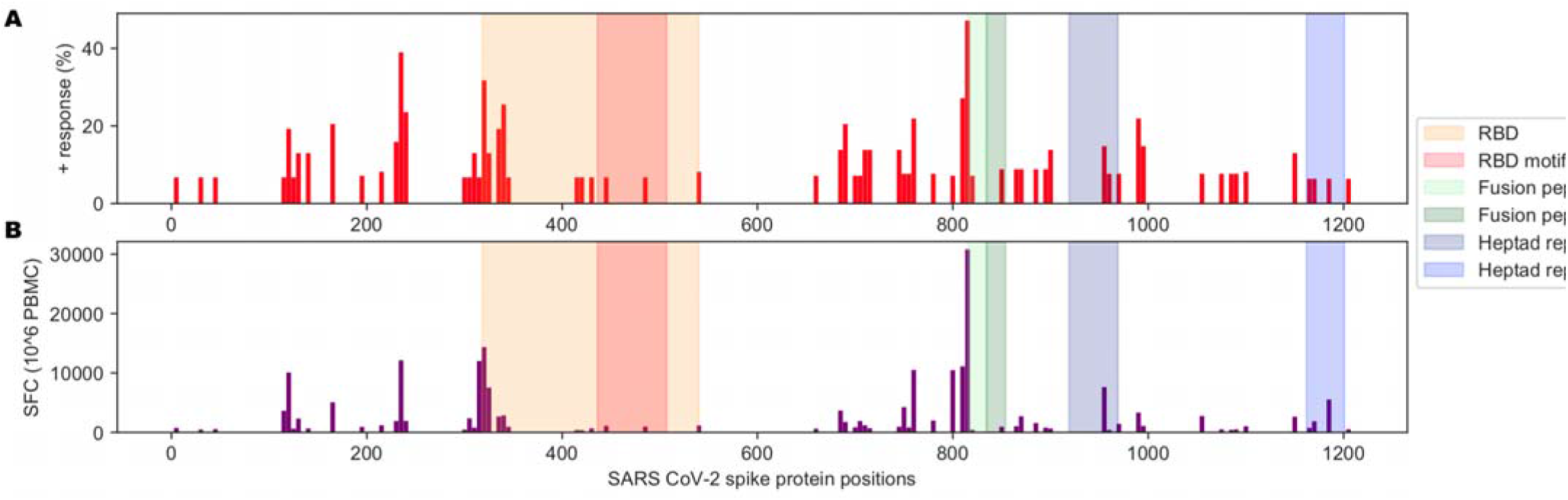
Immunological history of relevance to SARS-CoV-2. (A) Barplot showing the percentage of positive responses toward SARS-CoV-2 peptides from unexposed individuals. (B) Barplot showing the number of spot-forming cells (SFC) for tested SARS-CoV-2 peptides against PBMCs from unexposed individuals. Data from Table S1 from (46).

**Figure 5.**
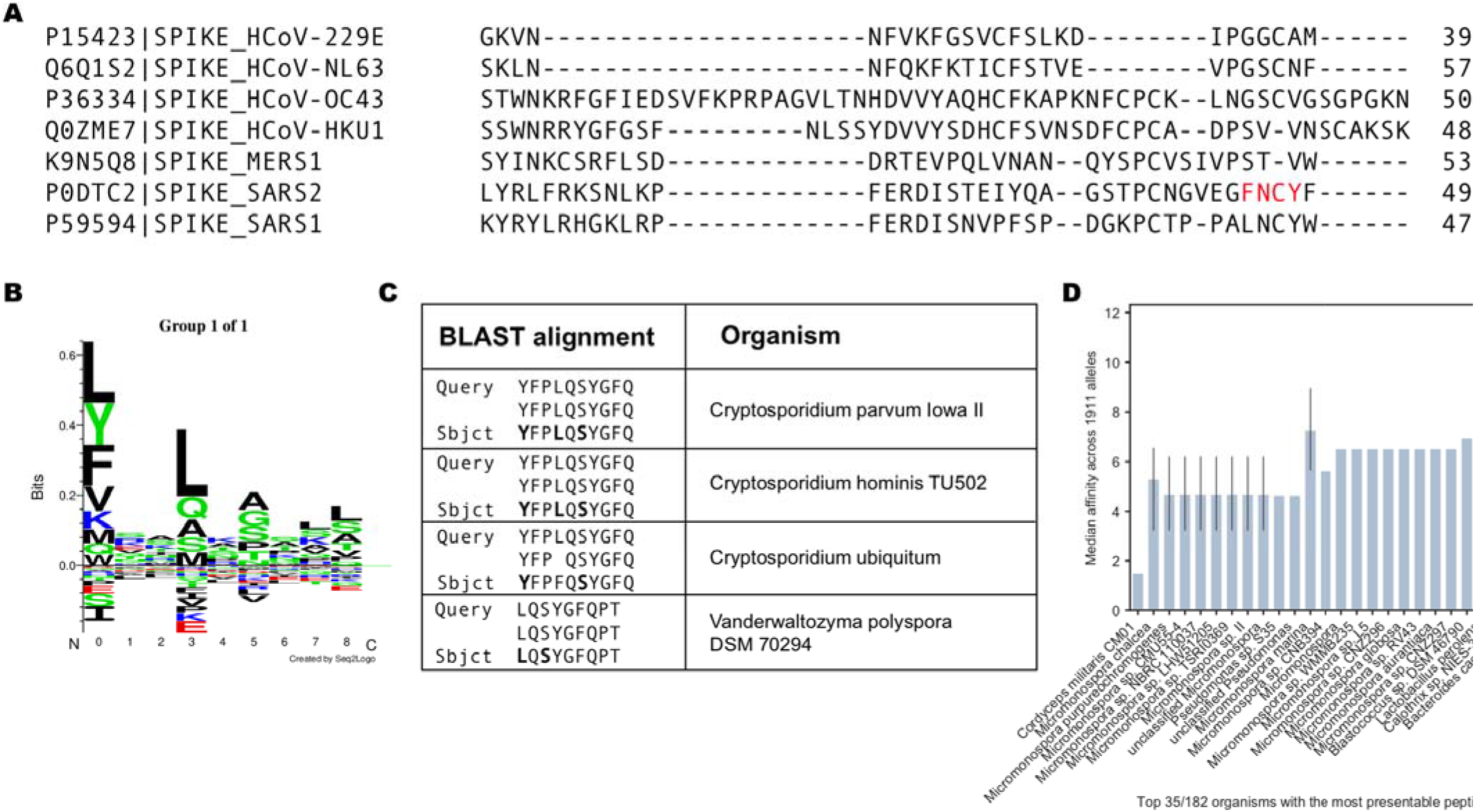
Learned immunity to other targets that could support T cell responses to SARS-CoV-2. (A) Multiple sequence alignment between SARS-CoV-2, SARS1, MERS, and other human coronaviruses, focusing on the region surrounding the FNCY B cell epitope. (B) SeqLogo plot obtained by clustering IEDB peptides reported to bind to DRB1*01:01. (C) Top results after blasting the FNCYFPLQSYGFQPT peptide against all reference proteins. (D) Barplot describing best peptide affinities across MHC-II alleles of the top 35 unique organisms with one or more peptides matching a peptide with high similarity to 15mers +/− 30aa from the FNCY binding epitope based on BLAST analysis. The closer to 0, the greater the binding potential.

A notable exception in Mateus’ results is peptide 486FNCYFPLQSYGFQPT500, which was reported to induce a CD4 T cell response in an unexposed individual. In this case, the peptide was restricted by HLA-DRB1*0101 or HLA-DQA1*0101/DQB1*0501. We found that the peptide sequence had greater *in silico* predicted affinity to HLA-DRB1*0101. To explain the conundrum, we blasted this peptide against the “refseq_protein” database excluding SARS-CoV-2 (Methods). Surprisingly, the sequences with the best homology for this query were not from coronaviruses but rather from common pathogens, first among them parasites of the *Cryptosporidium* genus of apicomplexan parasitic alveolates. These sequences included conserved anchor positions for the HLA-DRB*0101 allele making it plausible that a prior exposure could account for the formation of a memory CD4 T cell response (**Fig 5B-C**). To further assess the potential for other prior exposures in generating immune memory for sequences proximal to the FNCY patch we blasted all 15mers within +/− 30 amino acids of the FNCY patch and filtered the resulting sequences based on restriction by consensus MHC-II supertypes (50) (**Supplemental Table 2**). We found peptides associated with multiple microbial organisms that may meet the criteria to potentially generate CD4 T cell memory relevant to th RBM of SARS-CoV-2 (**Fig 5D**).

## Discussion

SARS-CoV-2 uses the RBD of the spike protein to bind to the ACE2 receptor on target cells. The actual contact with ACE2 is mediated by a discrete number of amino acids that have been visualized by cryo-EM (Lan et al., 2020; Shang et al., 2020). Although several SARS-related coronaviruses share 75% homology and interact with ACE2 on target cells (Ge et al., 2013; Ren et al., 2008; Yang et al., 2015) the RBM in SARS-CoV-2 is unique to this virus. *In vitro* binding measurements show that SARS-CoV-2 RBD binds to ACE2 with an affinity in the low nanomolar range (Walls et al., 2020). Mutations in this motif could be detrimental to the virus’s ability to infect ACE2 positive human cells. Since the RBD is an immunodominant site in the antibody response in humans (42) it is not surprising that the paratope of some antibodies isolated from convalescent individuals via single B cell VH/VL cloning, and selected on the basis of high neutralization potency, all seem to bind a surface encompassing the FNCY patch in the RBM (35,37–41,53). Arguably, this motif corresponds to a relevant B cell epitope in the spike protein of SARS-CoV-2 and is a logical target of potent neutralizing antibodies.

Although antibodies directed to this site have been isolated by different groups, little is known about their contribution to the pool of antibodies in serum of SARS-CoV-2 infected individuals, but evidence suggests they are likely to be rare. In one study they were found to represent a subdominant fraction of the anti-RBD response (41) while the estimated frequency of antigen-specific B cells ranges from 0.07 to 0.005% of all the total B cells in COVID-19 convalescent individuals (54). In a second study, the identification of two ultra-potent NAbs having a paratope involving the FNCY patch required screening of 800 clones from twelve individuals (53). This suggests that a potent NAb response to a mutationally constrained RBM epitope is a rare component of the total anti-virus response consistent, with the observation that there is no correlation between RBM site-specific neutralizing antibodies and serum half-maximal neutralization titer (NT50) (54). Here we show that the core RBM B cell epitope is apparently uncoupled from preferential T-B pairing, a prerequisite for a coordinated activation of B cells against the pathogen. We analyzed MHC-II binding of 15mer peptides in the spike protein upstream (-50 aa) or downstream (+50 aa) of the central RBM B cell epitope and found both low coverage by 1911 common MHC-II alleles and a depletion of binding 15mers proximal to the FNCY patch versus other exposed areas on the spike protein. This could be due to the fact that a sizeable proportion (40%) of CD4 T cells responding to the spike protein are memory responses found in SARS-CoV-2 unexposed individuals (44,55) or other structural protein of SARS-CoV-2 such as the N protein (45). Thus, it is possible that these conserved responses are used as a decoy mechanism to polarize the response away from the RBM. However, this does not rule out the contribution of a bias in frequency of specific B cells in the available repertoire.

Corroboration to our hypothesis also comes from Mateus *et al.* (46) who tested sixty-six 15mer peptides of the spike protein in SARS-CoV-2 unexposed individuals and found that CD4 T cell responses against this narrow RBM site account for only 2/110 (1.8%) of the total CD4 T cell response to 15mer peptides of the spike protein. Surprisingly, a CD4 T cell response against peptide FNCYFPLQSYGFQPT was by CD4 T cells of an unexposed individual. Since this peptide has low homology with previous human coronaviruses, we reasoned that this could either represent a case of TCR cross-reactivity since a single TCR can engage large numbers of unique MHC/peptide combinations without requiring degeneracy in their recognition (56,57). Remarkably, however, a BLAST analysis revealed a 10 amino acid sequence match with proteins from pathogens including those from the *Cryptosporidium* genus, with identity in binding motif and anchor residues (agretope) for the restricting MHC-II allele strongly suggesting peptide cross-reactivity. *Cryptosporidium hominis* is a parasite that causes watery diarrhea that can last up to 3 weeks in immunocompetent patients (58). Additional possibilities for cross-reactivity to the RBM, albeit of a lesser stringency, involve antigens from *Micromonospora*, *Pseudomonas, Blastococcus, Lactobacillus, and Bacteroides* (**Fig 5D**). Thus, it appears as if memory CD4 T cells reactive with peptides in the RBM may reflect the immunological history of the individual that, as evidenced by this case, can be unrelated to infection by other coronaviruses. Interestingly, the great majority (64-88%) of COVID-19 positive individuals in homeless shelters in Los Angeles and Boston were found to be asymptomatic (59). This suggests that the status of the immune system, which itself reflects past antigenic exposure, may be a determining factor in the generation of a protective immune response after SARS-CoV-2 infection.

The findings reported herein have considerable implications for natural immunity to SARS-CoV-2. The fact that there seems to be an overall suboptimal T-B preferential pairing suggests that B cells that respond to the RBM B cell epitope may receive inadequate T cell help. This is consistent with the observation that in general potent neutralizing antibodies to the RBM undergo very limited somatic mutation (38,53) and are by and large in quasi-germline configuration (60). Since T cell help is also necessary to initiate somatic hypermutation in B cell through CD40 or CD38 signaling in the germinal center (61), it follows that one important implication of our study is that defective T-B pairing may negatively influence the normal process of germinal center maturation of the B cell response in response to SARS-CoV-2 infection in a critical way.

Which antigens can generate T cell responses depends on the binding specificities of MHC-II molecules, which are highly polymorphic in the human population. We noted a general trend for MHC-II alleles to less effectively present peptides from the RBM region, but also observed some variability across MHC-II supertypes. The main DP and DP2 haplotypes were both common and had the highest potential to present peptides, suggesting that most individuals should carry at least one allele capable of presenting peptides in this region. Which of the two DP haplotypes was more common varied by ancestral population, thus it is possible that differences in the haplotypes could translate to differences in T-B cooperativity levels within groups, though binding affinities for epitopes near the FNCY patch were similar for both. DQ and DR supertypes were less able to present peptides near FNCY, with the exception of DR4, which is among the less common supertypes. Importantly, our analysis was limited to predicted affinity of peptides to MHC-II, and other characteristics such as expression levels, stability or differences in interactions with molecular chaperones likely also contribute to whether FNCY proximal peptides are available to support B-T cooperation (62).

In light of our findings, it can be predicted that, in general, a specific RBM antibody response may be short-lived and that residual immunity from a primary infection may not be sufficient to prevent reinfection after 6-9 months. Sporadic cases of re-infection have been reported by the media in Hong Kong and Nevada (63). A third case has been reported in a care-home resident who after the second infection produced only low levels of antibodies (64). Finally, silent re-infections in young workers in a COVID-19 ward who tested positive for the new coronavirus and became reinfected several months later with no symptoms in either instance have been reported (65). It is tempting to speculate that waning antibody levels or a poorly developed specific NAb antibody response to SARS-CoV-2 can potentially put people at risk of reinfection. Other factors to consider are a bias in the available B cell repertoire in the population and the extent to which a defective T-B cooperation influences the longevity of terminally differentiated plasma cells in the bone marrow (66).

In summary, we provide evidence that MHC-II constrains the CD4 T cell response for epitopes that are best positioned to facilitate T-B pairing in generating and sustaining a potent neutralizing antibody response against a mutationally constrained RBM B cell epitope. Furthermore, we show that the immunological history of the individual, not necessarily related to infection by other coronaviruses, may confer immunologic advantage. Finally, these findings may have implications for the quality and persistence of a protective, neutralizing antibody response to RBM induced by current SARS-CoV-2 vaccines.

## Materials and Methods

Data and code are available at https://github.com/cartercompbio/SARS_CoV_2_T-B_co-op.

### Affinity analysis

NetMHCIIpan version 4.0 was used to predict peptide-MHC-II affinity (69) for generated 15mers along the SARS-CoV-2 spike protein.

### Spike protein analyses

SARS-CoV-2 spike protein sequence and protein regions were obtained from https://www.uniprot.org/uniprot/P0DTC2. Glycan data were obtained from (49) and true-positive sites were aggregated across 3 replicates. To assess depletion of effective binders near the FNCY patch, we performed a Fisher’s exact test for binding (median affinity across common alleles <10) versus proximity (+/− 50 amino acids) to FNCY for positions free of glycans. We excluded positions within 10 amino acids of a glycan using the data obtained from Watanabe *et al*. and added a pseudocount of 1.

The SARS1, MERS1, HCoV-229E, HCoV-NL63, HCoV-OC43, and HCoV-HKU1 spike protein sequences were also downloaded from UniProt (P59594, K9N5Q8, P15423, Q6Q1S2, P36334, Q0ZME7, respectively). Multiple sequence alignment was performed on the EMBL-EBI Clustal Omega web server using default parameters (70).

### Structure analysis

The 6M0J 3D X-ray structure for the protein complex containing the SARS-CoV-2 spike protein RBD (P0DTC2) interaction with ACE2 (Q9BYF1) from (36). The structure figures were prepared using VMD (71).

### Supertype analysis

Supertypes were obtained from (50). All alpha/beta combinations spanning any of these types were included, resulting in 279 alleles. US supertype frequencies for alleles in DRB1 and DQB1 were obtained from the Be the Match registry (51), US frequencies for alleles in DPB1 were obtained from (52) as DPB1 was not available from the Be the Match registry. Available allele frequencies within each supertype were summed for Fig 3C.

### Motif analysis

All 13-20mer peptides adhering to the following parameters were downloaded from the IEDB (72): MHC-II assay, positive only, DRB1*01:01 allele, linear peptides; and any peptides with post-translational modifications or noncanonical amino acids were removed. The remaining 10,117 peptides were input into Gibbs cluster v2.0 (73) using the default MHC-II ligand parameters.

### BLAST analysis

15mers were generated along a sliding window +/− 30 amino acids from the FNCY patch start and end (455-518, 0-index) and input into NCBI BLAST (74) using the ‘refseq_protein’ database and excluding SARS-CoV-2 (taxid:2697049). Identified peptides (**Supplemental table 2**) were then evaluated for binding affinity and any peptide binding to at least one allele was retained for Fig 5D.

## Acknowledgements

This work was supported by an NIH National Library of Medicine Training Grant T15LM011271 to A.C., an Emerging Leader Award from The Mark Foundation for Cancer Research, grant #18-022-ELA and a CIFAR fellowship to H.C. and RO1 CA220009 to M.Z. and H.C. The graphical abstract was created using BioRender and used the PDB (67) structure 6VXX from (68).

## Author Contributions

Original concept, M.Z.; project supervision, H.C. and M.Z.; project planning and experimental design, A.C., M.Z., and H.C.; data acquisition, processing, and analysis, A.C. and K.O.; preparation of paper, A.C., M.Z., and H.C.

## Supplemental Table Legends

Supplemental Table 1: SARS-CoV-2 neutralizing antibody residues and references used to generate Fig 1D.

Supplemental Table 2: BLAST-identified peptides with affinity, and binding fraction.

## References

1. Wölfel R, Corman VM, Guggemos W, Seilmaier M, Zange S, Müller MA, et al. Virological assessment of hospitalized patients with COVID-2019. Nature. 2020 May;581(7809):465–9.

2. Long Q-X, Tang X-J, Shi Q-L, Li Q, Deng H-J, Yuan J, et al. Clinical and immunological assessment of asymptomatic SARS-CoV-2 infections. Nat Med. 2020 Aug;26(8):1200–4.

3. Rydyznski Moderbacher C, Ramirez SI, Dan JM, Grifoni A, Hastie KM, Weiskopf D, et al. Antigen-Specific Adaptive Immunity to SARS-CoV-2 in Acute COVID-19 and Associations with Age and Disease Severity. Cell [Internet]. 2020 Sep 16; Available from: http://dx.doi.org/10.1016/j.cell.2020.09.038

4. Prévost J, Gasser R, Beaudoin-Bussières G, Richard J, Duerr R, Laumaea A, et al. Cross-sectional evaluation of humoral responses against SARS-CoV-2 Spike. Cell Rep Med. 2020 Sep 30;100126.

5. Mitchison NA. T-cell-B-cell cooperation. Nat Rev Immunol. 2004 Apr;4(4):308–12.

6. Jacob J, Kelsoe G, Rajewsky K, Weiss U. Intraclonal generation of antibody mutants in germinal centres. Nature. 1991 Dec 5;354(6352):389–92.

7. Berek C, Berger A, Apel M. Maturation of the immune response in germinal centers. Cell. 1991 Dec 20;67(6):1121–9.

8. Zanetti M, Glotz D. Considerations on thymus-dependent and -independent antigens in acquired and natural immunity. Ann Inst Pasteur Immunol. 1988 Mar;139(2):192–3.

9. Claman HN, Chaperon EA, Triplett RF. Thymus-marrow cell combinations. Synergism in antibody production. Proc Soc Exp Biol Med. 1966 Aug;122(4):1167–71.

10. Mitchison NA. The carrier effect in the secondary response to hapten-protein conjugates. I. Measurement of the effect with transferred cells and objections to the local environment hypothesis. Eur J Immunol. 1971 Jan;1(1):10–7.

11. Rajewsky K, Rottländer E, Peltre G, Müller B. The immune response to a hybrid protein molecule; specificity of secondary stimulation and of tolerance induction. J Exp Med. 1967 Oct 1;126(4):581–606.

12. Katz DH, Hamaoka T, Dorf ME, Benacerraf B. Cell interactions between histoincompatible T and B lymphocytes. The H-2 gene complex determines successful physiologic lymphocyte interactions. Proc Natl Acad Sci U S A. 1973 Sep;70(9):2624–8.

13. Sprent J. Restricted helper function of F1 hybrid T cells positively selected to heterologous erythrocytes in irradiated parental strain mice. II. Evidence for restrictions affecting helper cell induction and T-B collaboration, both mapping to the K-end of the H-2 complex. J Exp Med. 1978 Apr 1;147(4):1159–74.

14. Jones B, Janeway CA Jr. Cooperative interaction of B lymphocytes with antigen-specific helper T lymphocytes is MHC restricted. Nature. 1981 Aug 6;292(5823):547–9.

15. Mitchison NA. The carrier effect in the secondary response to hapten-protein conjugates. II. Cellular cooperation. Eur J Immunol. 1971;1(1):18–27.

16. Janeway CA Jr. Cellular cooperation during in vivo anti-hapten antibody responses. I. The effect of cell number on the response. J Immunol. 1975 Apr;114(4):1394–401.

17. Shulman Z, Gitlin AD, Targ S, Jankovic M, Pasqual G, Nussenzweig MC, et al. T follicular helper cell dynamics in germinal centers. Science. 2013 Aug 9;341(6146):673–7.

18. Celada F, Sercarz EE. Preferential pairing of T-B specificities in the same antigen: the concept of directional help. Vaccine. 1988 Apr;6(2):94–8.

19. Manca F, Kunkl A, Fenoglio D, Fowler A, Sercarz E, Celada F. Constraints in T-B cooperation related to epitope topology on E. coli β-galactosidase. I. The fine specificity of T cells dictates the fine specificity of antibodies directed to conformation-dependent determinants. Eur J Immunol. 1985;15(4):345–50.

20. Bretscher P, Cohn M. A theory of self-nonself discrimination. Science. 1970 Sep 11;169(3950):1042–9.

21. Lanzavecchia A. Antigen-specific interaction between T and B cells [Internet]. Vol. 314, Nature. 1985. p. 537–9. Available from: http://dx.doi.org/10.1038/314537a0

22. Kroeger DR, Rudulier CD, Bretscher PA. Antigen presenting B cells facilitate CD4 T cell cooperation resulting in enhanced generation of effector and memory CD4 T cells. PLoS One. 2013 Oct 14;8(10):e77346.

23. Cassell D, Forman J. Linked recognition of helper and cytotoxic antigenic determinants for the generation of cytotoxic T lymphocytes. Ann N Y Acad Sci. 1988;532:51–60.

24. Gerloni M, Xiong S, Mukerjee S, Schoenberger SP, Croft M, Zanetti M. Functional cooperation between T helper cell determinants. Proc Natl Acad Sci U S A. 2000 Nov 21;97(24):13269–74.

25. Berzofsky JA, Richman LK, Killion DJ. Distinct H-2-linked Ir genes control both antibody and T cell responses to different determinants on the same antigen, myoglobin. Proc Natl Acad Sci U S A. 1979 Aug;76(8):4046–50.

26. Berzofsky JA, Schechter AN, Shearer GM, Sachs DH. Genetic control of the immune response to staphylococcal nuclease. III. Time-course and correlation between the response to native nuclease and the response to its polypeptide fragments. J Exp Med. 1977 Jan 1;145(1):111–22.

27. Berzofsky JA, Schechter AN, Shearer GM, Sachs DH. Genetic control of the immune response to staphylococcal nuclease. IV. H-2-linked control of the relative proportions of antibodies produced to different determinants of native nuclease. J Exp Med. 1977 Jan 1;145(1):123–35.

28. Zanetti M, Sercarz E, Salk J. The immunology of new generation vaccines. Immunol Today. 1987;8(1):18–25.

29. Celada F, Kunkl A, Manca F, Fenoglio D, Fowler A, Krzych U, et al. Preferential pairings in T-B encounters utilizing Th cells directed against discrete portions of b-galactosidase and B cells primed with the native enzyme or a hapten epitope. Regulation of the Immune System. 1984;637–46.

30. Sette A, Moutaftsi M, Moyron-Quiroz J, McCausland MM, Davies DH, Johnston RJ, et al. Selective CD4+ T cell help for antibody responses to a large viral pathogen: deterministic linkage of specificities. Immunity. 2008 Jun;28(6):847–58.

31. Lv Z, Deng Y-Q, Ye Q, Cao L, Sun C-Y, Fan C, et al. Structural basis for neutralization of SARS-CoV-2 and SARS-CoV by a potent therapeutic antibody. Science. 2020 Sep 18;369(6510):1505–9.

32. Pinto D, Park Y-J, Beltramello M, Walls AC, Tortorici MA, Bianchi S, et al. Cross-neutralization of SARS-CoV-2 by a human monoclonal SARS-CoV antibody. Nature. 2020 Jul;583(7815):290–5.

33. Yuan M, Wu NC, Zhu X, Lee C-CD, So RTY, Lv H, et al. A highly conserved cryptic epitope in the receptor binding domains of SARS-CoV-2 and SARS-CoV. Science. 2020 May 8;368(6491):630–3.

34. Piccoli L, Park Y-J, Tortorici MA, Czudnochowski N, Walls AC, Beltramello M, et al. Mapping Neutralizing and Immunodominant Sites on the SARS-CoV-2 Spike Receptor-Binding Domain by Structure-Guided High-Resolution Serology. Cell [Internet]. 2020 Sep 16; Available from: http://dx.doi.org/10.1016/j.cell.2020.09.037

35. Barnes CO, West AP Jr, Huey-Tubman KE, Hoffmann MAG, Sharaf NG, Hoffman PR, et al. Structures of Human Antibodies Bound to SARS-CoV-2 Spike Reveal Common Epitopes and Recurrent Features of Antibodies. Cell. 2020 Aug 20;182(4):828–42.e16.

36. Lan J, Ge J, Yu J, Shan S, Zhou H, Fan S, et al. Structure of the SARS-CoV-2 spike receptor-binding domain bound to the ACE2 receptor. Nature. 2020 May;581(7807):215–20.

37. Liu L, Wang P, Nair MS, Yu J, Rapp M, Wang Q, et al. Potent neutralizing antibodies against multiple epitopes on SARS-CoV-2 spike. Nature. 2020 Aug;584(7821):450–6.

38. Rogers TF, Zhao F, Huang D, Beutler N, Burns A, He W-T, et al. Isolation of potent SARS-CoV-2 neutralizing antibodies and protection from disease in a small animal model. Science. 2020 Aug 21;369(6506):956–63.

39. Shi R, Shan C, Duan X, Chen Z, Liu P, Song J, et al. A human neutralizing antibody targets the receptor-binding site of SARS-CoV-2. Nature. 2020 Aug;584(7819):120–4.

40. Wu Y, Wang F, Shen C, Peng W, Li D, Zhao C, et al. A noncompeting pair of human neutralizing antibodies block COVID-19 virus binding to its receptor ACE2. Science. 2020 Jun 12;368(6496):1274–8.

41. Zost SJ, Gilchuk P, Case JB, Binshtein E, Chen RE, Nkolola JP, et al. Potently neutralizing and protective human antibodies against SARS-CoV-2. Nature. 2020 Aug;584(7821):443–9.

42. Premkumar L, Segovia-Chumbez B, Jadi R, Martinez DR, Raut R, Markmann A, et al. The receptor binding domain of the viral spike protein is an immunodominant and highly specific target of antibodies in SARS-CoV-2 patients. Sci Immunol [Internet]. 2020 Jun 11;5(48). Available from: http://dx.doi.org/10.1126/sciimmunol.abc8413

43. Starr TN, Greaney AJ, Hilton SK, Ellis D, Crawford KHD, Dingens AS, et al. Deep Mutational Scanning of SARS-CoV-2 Receptor Binding Domain Reveals Constraints on Folding and ACE2 Binding. Cell. 2020 Sep 3;182(5):1295–310.e20.

44. Grifoni A, Weiskopf D, Ramirez SI, Mateus J, Dan JM, Moderbacher CR, et al. Targets of T Cell Responses to SARS-CoV-2 Coronavirus in Humans with COVID-19 Disease and Unexposed Individuals. Cell. 2020 Jun 25;181(7):1489–501.e15.

45. Le Bert N, Tan AT, Kunasegaran K, Tham CYL, Hafezi M, Chia A, et al. SARS-CoV-2-specific T cell immunity in cases of COVID-19 and SARS, and uninfected controls. Nature. 2020 Aug;584(7821):457–62.

46. Mateus J, Grifoni A, Tarke A, Sidney J, Ramirez SI, Dan JM, et al. Selective and cross-reactive SARS-CoV-2 T cell epitopes in unexposed humans. Science [Internet]. 2020 Aug 4; Available from: http://dx.doi.org/10.1126/science.abd3871

47. Yuan M, Liu H, Wu NC, Lee C-CD, Zhu X, Zhao F, et al. Structural basis of a shared antibody response to SARS-CoV-2. Science. 2020 Aug 28;369(6507):1119–23.

48. Dosset M, Castro A, Carter H, Zanetti M. Telomerase and CD4 T Cell Immunity in Cancer. Cancers [Internet]. 2020 Jun 25;12(6). Available from: http://dx.doi.org/10.3390/cancers12061687

49. Watanabe Y, Allen JD, Wrapp D, McLellan JS, Crispin M. Site-specific glycan analysis of the SARS-CoV-2 spike. Science. 2020 Jul 17;369(6501):330–3.

50. Greenbaum J, Sidney J, Chung J, Brander C, Peters B, Sette A. Functional classification of class II human leukocyte antigen (HLA) molecules reveals seven different supertypes and a surprising degree of repertoire sharing across supertypes. Immunogenetics. 2011 Jun;63(6):325–35.

51. Maiers M, Gragert L, Klitz W. High-resolution HLA alleles and haplotypes in the United States population. Hum Immunol. 2007 Sep;68(9):779–88.

52. Du Z. HLA-DPA1 and HLA-DPB1 Frequencies in the US Populations [Internet]. 2017 American Transplant Congress; 2017 Apr 30 [cited 2020 Sep 30]; Chicago, IL. Available from: https://atcmeetingabstracts.com/abstract/hla-dpa1-and-hla-dpb1-frequencies-in-the-us-populations/

53. Tortorici MA, Beltramello M, Lempp FA, Pinto D, Dang HV, Rosen LE, et al. Ultrapotent human antibodies protect against SARS-CoV-2 challenge via multiple mechanisms. Science [Internet]. 2020 Sep 24; Available from: http://dx.doi.org/10.1126/science.abe3354

54. Robbiani DF, Gaebler C, Muecksch F, Lorenzi JCC, Wang Z, Cho A, et al. Convergent antibody responses to SARS-CoV-2 in convalescent individuals. Nature. 2020 Aug;584(7821):437–42.

55. Braun J, Loyal L, Frentsch M, Wendisch D, Georg P, Kurth F, et al. SARS-CoV-2-reactive T cells in healthy donors and patients with COVID-19. Nature [Internet]. 2020 Jul 29; Available from: http://dx.doi.org/10.1038/s41586-020-2598-9

56. Birnbaum ME, Mendoza JL, Sethi DK, Dong S, Glanville J, Dobbins J, et al. Deconstructing the peptide-MHC specificity of T cell recognition. Cell. 2014 May 22;157(5):1073–87.

57. Selin LK, Cornberg M, Brehm MA, Kim S-K, Calcagno C, Ghersi D, et al. CD8 memory T cells: cross-reactivity and heterologous immunity. Semin Immunol. 2004 Oct;16(5):335–47.

58. Gharpure R, Perez A, Miller AD, Wikswo ME, Silver R, Hlavsa MC. Cryptosporidiosis Outbreaks - United States, 2009-2017. MMWR Morb Mortal Wkly Rep. 2019 Jun 28;68(25):568–72.

59. Oran DP, Topol EJ. Prevalence of Asymptomatic SARS-CoV-2 Infection : A Narrative Review. Ann Intern Med. 2020 Sep 1;173(5):362–7.

60. Kreer C, Zehner M, Weber T, Ercanoglu MS, Gieselmann L, Rohde C, et al. Longitudinal Isolation of Potent Near-Germline SARS-CoV-2-Neutralizing Antibodies from COVID-19 Patients. Cell. 2020 Sep 17;182(6):1663–73.

61. Bergthorsdottir S, Gallagher A, Jainandunsing S, Cockayne D, Sutton J, Leanderson T, et al. Signals that initiate somatic hypermutation of B cells in vitro. J Immunol. 2001 Feb 15;166(4):2228–34.

62. Anczurowski M, Hirano N. Mechanisms of HLA-DP Antigen Processing and Presentation Revisited. Trends Immunol. 2018 Dec;39(12):960–4.

63. Tillett RL, Sevinsky JR, Hartley PD, Kerwin H, Crawford N, Gorzalski A, et al. Genomic evidence for reinfection with SARS-CoV-2: a case study. Lancet Infect Dis [Internet]. 2020 Oct 12; Available from: http://www.sciencedirect.com/science/article/pii/S1473309920307647

64. Goldman JD, Wang K, Roltgen K, Nielsen SCA, Roach JC, Naccache SN, et al. Reinfection with SARS-CoV-2 and Failure of Humoral Immunity: a case report. medRxiv [Internet]. 2020 Sep 25; Available from: http://dx.doi.org/10.1101/2020.09.22.20192443

65. Gupta V, Bhoyar RC, Jain A, Srivastava S, Upadhayay R, Imran M, et al. Asymptomatic reinfection in two healthcare workers from India with genetically distinct SARS-CoV-2. Clin Infect Dis [Internet]. 2020 Sep 23; Available from: http://dx.doi.org/10.1093/cid/ciaa1451

66. Slifka MK, Matloubian M, Ahmed R. Bone marrow is a major site of long-term antibody production after acute viral infection. J Virol. 1995 Mar;69(3):1895–902.

67. Sehnal D, Rose AS, Koča J, Burley SK, Velankar S. Mol*: towards a common library and tools for web molecular graphics. In: MolVa: Workshop on Molecular Graphics and Visual Analysis of Molecular Data, Brno, Czech Republic Eurographics. 2018.

68. Walls AC, Park Y-J, Tortorici MA, Wall A, McGuire AT, Veesler D. Structure, Function, and Antigenicity of the SARS-CoV-2 Spike Glycoprotein. Cell. 2020 Apr 16;181(2):281–92.e6.

69. Reynisson B, Alvarez B, Paul S, Peters B, Nielsen M. NetMHCpan-4.1 and NetMHCIIpan-4.0: improved predictions of MHC antigen presentation by concurrent motif deconvolution and integration of MS MHC eluted ligand data. Nucleic Acids Res [Internet]. 2020; Available from: https://academic.oup.com/nar/advance-article-abstract/doi/10.1093/nar/gkaa379/5837056

70. Madeira F, Park YM, Lee J, Buso N, Gur T, Madhusoodanan N, et al. The EMBL-EBI search and sequence analysis tools APIs in 2019. Nucleic Acids Res. 2019 Jul 2;47(W1):W636–41.

71. Humphrey W, Dalke A, Schulten K. VMD: visual molecular dynamics. J Mol Graph. 1996 Feb;14(1):33–8, 27–8.

72. Vita R, Mahajan S, Overton JA, Dhanda SK, Martini S, Cantrell JR, et al. The Immune Epitope Database (IEDB): 2018 update. Nucleic Acids Res. 2019 Jan 8;47(D1):D339–43.

73. Andreatta M, Lund O, Nielsen M. Simultaneous alignment and clustering of peptide data using a Gibbs sampling approach. Bioinformatics. 2013 Jan 1;29(1):8–14.

74. Sayers EW, Barrett T, Benson DA, Bolton E, Bryant SH, Canese K, et al. Database resources of the National Center for Biotechnology Information. Nucleic Acids Res. 2011 Jan;39(Database issue):D38–51.

