## Supplementary figures for "MHC-II constrains the natural neutralizing antibody response to the SARS-CoV-2 spike RBM in humans"

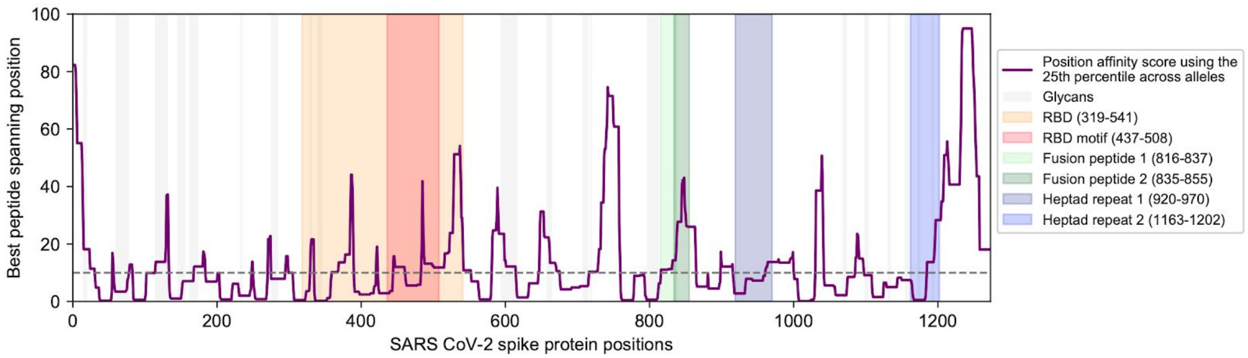


Supplemental Figure 1. Distribution of position scores along the spike protein using the 25th percentile affinity instead of the median affinity.

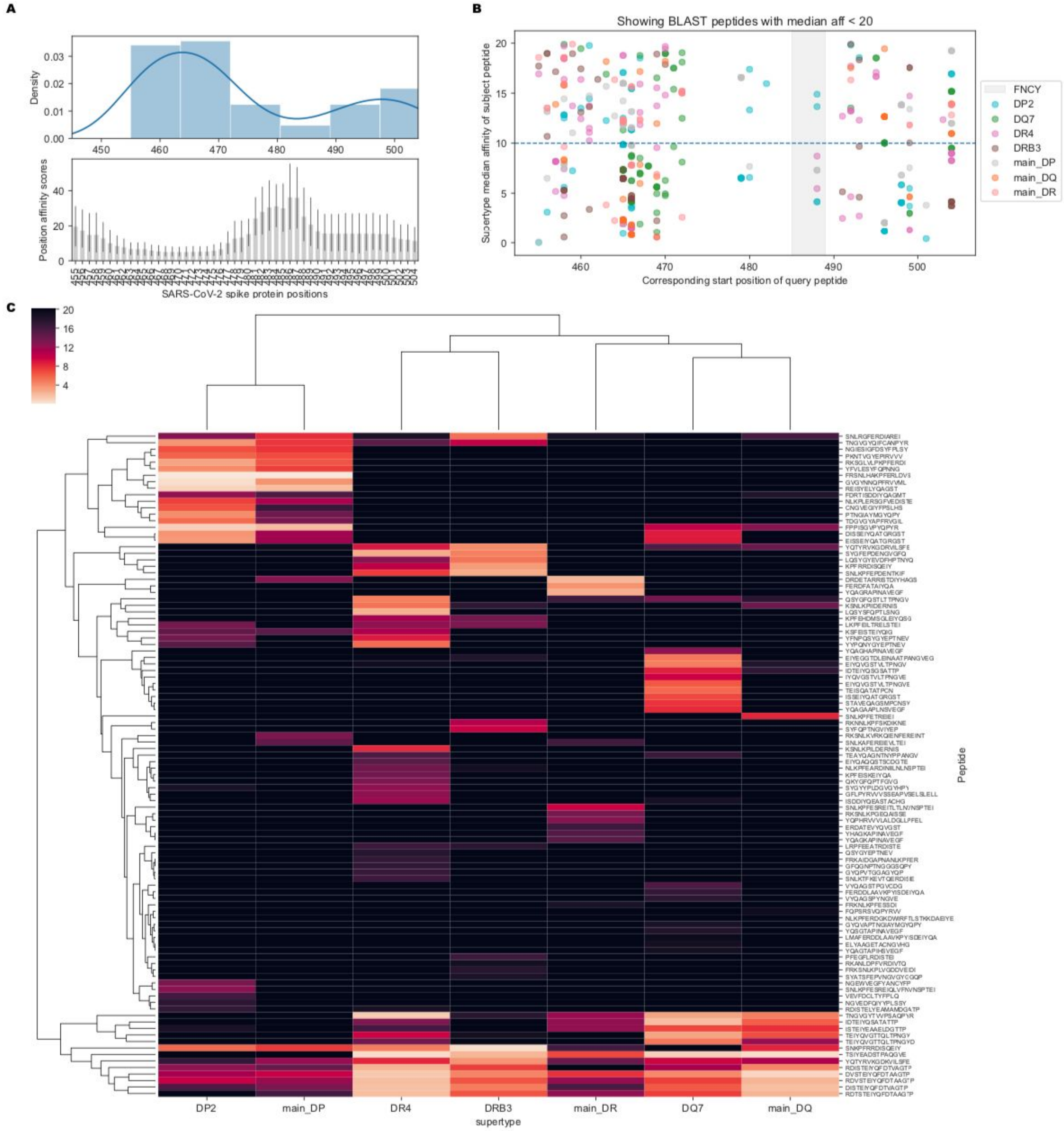


Supplemental Figure 2. Overview of subject peptides that bind at least one retrieved from BLAST search. (A) Pileup of corresponding query peptides' start positions of BLAST-identified peptides that bind to at least one common MHC-II allele. The below barplot shows Figure 3A for reference: the aggregated position scores across supertypes for positions proximal to FNCY. (B) Scatterplot showing the median supertype affinities of BLAST-identified peptides that may bind (median affinity <20) along the corresponding start positions of queried peptides along the spike protein. The FNCY motif region is highlighted in grey. (C) Clustermap showing the median supertype affinities of BLAST-identified peptides that may bind (median affinity <20) to at least one supertype. Median affinities greater than 20 have been adjusted to 20 for better visualization of binding peptides.
